# Systematic selection of reference genes for normalization of circulating RNA transcripts in pregnant women based on RNA-seq data

**DOI:** 10.1101/165654

**Authors:** Stephen S. C. Chim, Karen K. W. Wong, Claire Y. L. Chung, Stephanie K. W. Lam, Jamie S. L. Kwok, Chit-Ying Lai, Yvonne K. Y. Cheng, Annie S. Y. Hui, Meng Meng, Oi-Ka Chan, Stephen K. W. Tsui, Keun-Young Lee, Ting-Fung Chan, Tak-Yeung Leung

**Affiliations:** Department of Obstetrics & Gynaecology, Faculty of Medicine, The Chinese University of Hong Kong, Shatin, New Territories, Hong Kong SAR, China; (K.K.W.W.), (S.K.W.L.), (C.-Y.L.), (Y.K.Y.C.), (A.S.Y.H.), (M.M.), (O.K.C.); School of Life Sciences, Faculty of Science, The Chinese University of Hong Kong, Shatin, New Territories, Hong Kong SAR, China; (C.Y.L.C.), (T.-F.C.); Hong Kong Bioinformatics Center, The Chinese University of Hong Kong, Shatin, New Territories, Hong Kong SAR, China.; School of Biomedical Sciences, Faculty of Medicine, The Chinese University of Hong Kong, Shatin, New Territories, Hong Kong SAR, China; (J.S.L.K.), (S.K.W.T.); Department of Obstetrics & Gynaecology, Division of Maternal and Fetal Medicine, Kangnam Sacred Heart Hospital, College of Medicine, Hallym University, Seoul, Republic of Korea

**Keywords:** reference target, qRT-PCR, normalization, transcriptomics, blood biomarkers, geNorm, NormFinder, Removal of Unwanted Variation RUVSeq, technical variation, denoise

## Abstract

RNA transcripts circulating in peripheral blood represent an important source of non-invasive biomarkers. To accurately quantify the levels of a circulating transcript, one needs to normalize the data with internal control reference genes, which are detected at relatively constant levels across different blood samples. A few stably-expressed reference gene candidates have to be selected from transcriptome data before validation of their stable expression by reverse-transcription quantitative polymerase chain reaction. However, there is a lack of transcriptome, let alone whole-transcriptome, data from maternal blood. To overcome this shortfall, we performed RNA-seq on blood samples from women presented with preterm labor. Of 11215 exons detected in the maternal blood whole-transcriptome, we systematically identified a panel of 395 genes comprising exons that were detected at a *coefficient of variation* (*CV*) ranging from 7.75%-17.7%. Their levels were considerably less variable than any *GAPDH* exon (minimum *CV*, 27.3%). Upon validation, selected genes from this panel remained as more stably expressed than *GAPDH* in maternal blood. This panel is over-represented with genes involved with actin cytoskeleton, macromolecular complex and the integrin signaling pathway. This groundwork provides a starting point for systematically selecting reference gene candidates for normalizing the levels of circulating RNA transcripts in maternal blood.

## 1. Introduction

Quantitative polymerase chain reaction (qPCR) is a standard method for quantification of nucleic acid sequences [1,2]. Combined with a prior step of reverse transcription (RT), RT-qPCR has become a well-established technique to quantify the level of any mRNA transcript in a sample [3]. To control for experimental error between samples that can be introduced at a number of stages throughout the procedure, normalization of the RT-qPCR data is essential before analysis. This is usually achieved by the use of a reference gene as an internal control that is presumed to remain relatively constant across different samples. However, ideal reference genes rarely exist and finding suitable ones is not a trivial task. It has been demonstrated that a proper choice of reference genes is highly dependent on the tissues or cells under investigation [4]. Further, reference genes are highly specific for a particular experimental model, and validation for each situation, on an individual basis, is a crucial requirement [5].

Preterm birth (delivery before 37 weeks of gestation) is a major cause of neonatal morbidity and mortality [6]. Less than half of pregnant women presented with preterm labor end in spontaneous preterm birth (sPTB), while the remaining end in term birth (TB) on or after 37 weeks. To better understand sPTB, using gene expression microarrays, we systematically identified panels of RNA transcripts that are aberrantly expressed in the placentas [7] and maternal blood cell samples [8] collected from pregnancies undergoing sPTB. Since these preterm birth-associated transcripts are detectable in maternal whole blood, in this study, we aimed to identify reference genes suitable for normalizing RNA transcripts in the whole blood of women undergoing preterm labor.

Reference genes for normalizing RNA transcripts in the human circulatory system have been reported on patients with tuberculosis [5], schizophrenia and bipolar disorder [9] and cohort of healthy male and female adults [10]. Nevertheless, there is a lack of similar data on pregnant women. Based on a meta-analysis of publicly available gene expression microarray datasets on 1 053 blood samples, Cheng and colleagues have identified a panel of candidate reference genes for normalizing RT-qPCR data from peripheral blood across healthy, non-pregnant, individuals and patients with cancer or other abnormality [11]. Yet, whether those reference genes are suitable for normalization of data from pregnant women is unknown.

To address this paucity of relevant study, we embarked on a search for reference genes suitable for normalization of RT-qPCR data on whole blood collected from women during their presentation of preterm labor. Similar search for reference genes often relies on a candidate gene approach or gene expression microarray data, which is sometimes referred as transcriptome data. Such microarray-based transcriptome studies measure RNA levels based on oligonucleotide probes, which requires the prior knowledge of RNA transcript sequences, and thus are limited essentially to the more characterized genes of known sequences. To maximize the chance of finding suitable reference genes, in this study, we expanded this search to the whole-transcriptome dataset generated by RNA-sequencing (RNA-seq), which is not limited by probes of well-characterized gene sequences. We hypothesized that the whole-transcriptome of maternal blood harbors many RNA transcripts that are expressed at relatively constant levels. Based on RNA-seq dataset, we systematically identified 395 genes, comprising 458 exons, which were detected at a low variation across all tested maternal blood samples. Subsequently, using RT-qPCR, we assessed the expression stability of selected genes in another set of maternal blood samples. Our data suggest that the whole-transcriptome harbors gene candidates of higher expression stability than those commonly used reference genes.

In this study, we interpreted the RNA-seq data at the finer resolution of exons, which are sub-regions within an expressed gene transcript. We observed that different exons of the same gene were detected at considerably different levels of variation. Based on the exon-level RNA-seq data, one could readily pinpoint on the exon with the least variation in the design of RT-qPCR assays for the quantification of candidate reference genes. Further, we explored on how our list of 458 stably detected exons could be used as a starting point for the systematic identification of reference genes in other blood compartments and different group of patients. Lastly, we discussed the pros and cons of our exon-level RNA-seq approach and how it may be further improved for identifying reference genes.

## 2. Results

### 2.1. Whole-transcriptome profiling of maternal blood samples

#### 2.1.1. Overall results of the RNA-seq experiment

We obtained the informed consent and collected peripheral whole blood samples from pregnant women during their presentation of preterm labor. Twenty blood samples were subjected to strand-specific RNA-seq (ssRNA-seq). Forty libraries (2 technical replicates per blood sample) were constructed for strand-specific pair-end cDNA sequencing on the HiSeq 4000 sequencer (Illumina). We filtered out low-quality sequences, trimmed away adapter sequences (Trimmomatic, v.0.33) [12] and aligned the reads to the reference human genome (GRCh38, GENCODE release 23 primary assembly; aligner, STAR (v2.4.2) [13]). After all filtering and mapping, we rejected two libraries with high percentage of poor or unaligned reads and removed them from further analysis. The mean number of raw reads was 159 million per sample. Of the mean 93.6 million high-quality reads, 69.2% mapped to a unique location in the reference human genome. No remarkable difference was observed across the technical replicates.

#### 2.1.2. Identification of exons that are stably detected in maternal blood

Each gene comprises one or more sub-regions called exons. At the above sequencing depth, RNA-seq allows investigators to profile RNA levels not only at gene-level but also at the higher resolution of exon-level. Since the higher resolution exon-level data are advantageous for developing RT-qPCR assays, we summarized the counts of the reads mapped to each exon in GRCh38. To account for technical variations including the differing sequencing depth and amount of RNA input for each library, we normalized the count data using the Removal of Unwanted Variation (RUV) method, which is based on the factor analysis of control samples, such as technical replicates [14]. Unless otherwise stated, all further instances of “counts” in this paper refer to normalized counts. Of the 11 215 exons observed in our RNA-seq data, 4 579 exons were observed with at least one count in all libraries and were considered as robustly detected. For each exon, its RNA levels were calculated by the normalized counts of reads that mapped to that particular exonic sequence. Its mean RNA levels and coefficient of variation (*CV* = *standard deviation* divided by the *mean*) across all maternal blood samples were calculated. Figure 1 shows the *mean* and *CV* of the robustly detected exons in the *log_10_* scale.

**Figure 1.**
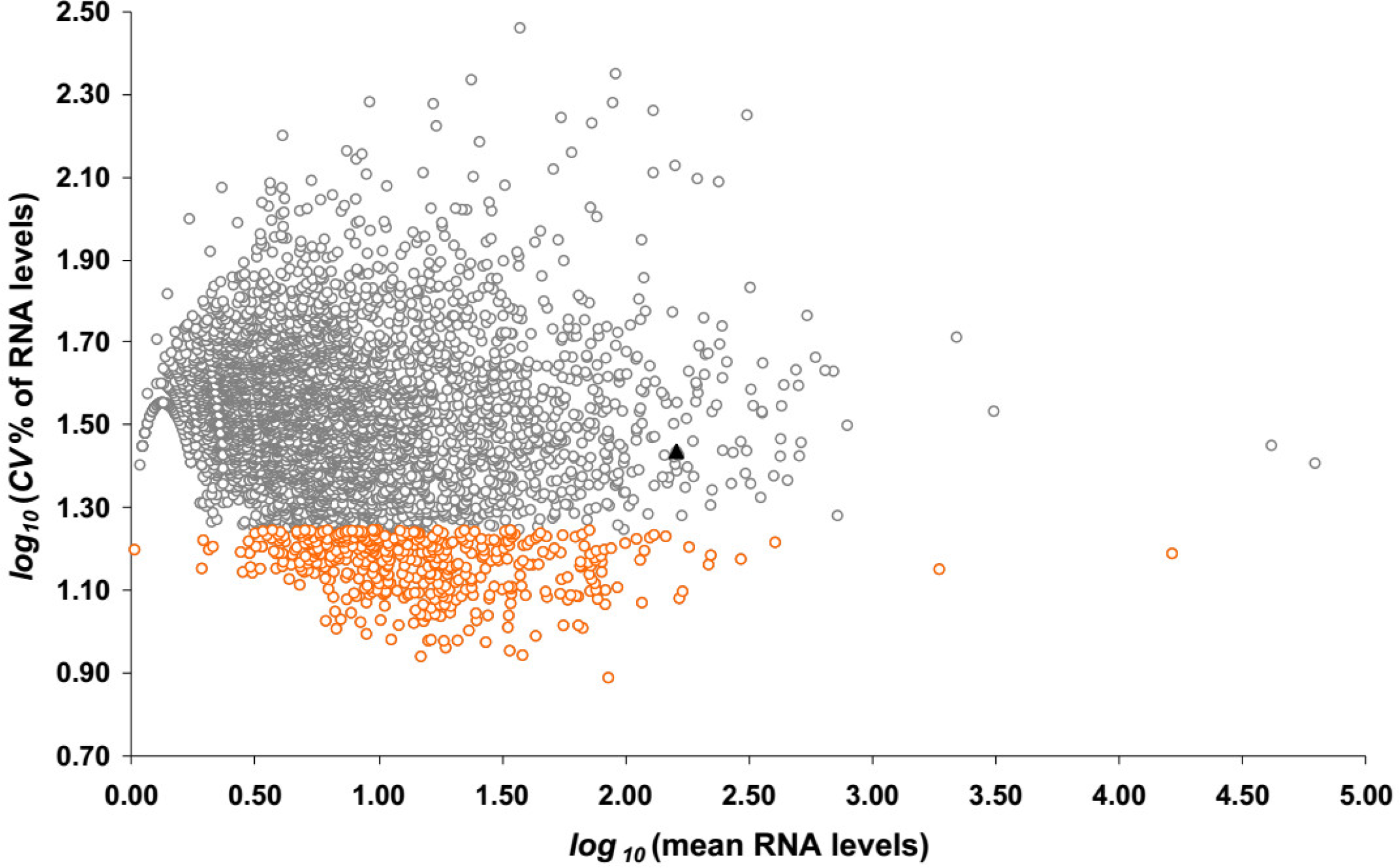
The *CV*% and *mean* of RNA expression levels of gene exons across all maternal blood samples based on RNA-seq. Each datapoint represents an exon. Shown are 4 579 exons that are robustly detected (minimum RNA level across all samples > 0 count). RNA levels are calculated by normalized read counts mapping to each exon. Highlighted in orange circles are 458 exons with *CV*% in the lowest 10^th^ percentile. Black triangle shows the data from the *GAPDH* transcript.

Without presuming normal distribution of the linear-scale data on RNA levels across the 4 579 exons, we describe the data as follows: The *median* (*interquartile range*, *IQR*) of the mean RNA levels was 5.7 counts (3.2 counts – 13 counts) and the *median* of the *CV* (*IQR*) of the RNA levels was 32.5% (24.0% to 42.0%). The *10^th^* and *90^th^ percentiles* of the *CV* are 17.7% and 53.8%, respectively. The Spearman rank order correlation coefficient was only -0.06 (*p*-value, 2 × 10^-4^) between the *mean* and the *CV* of the RNA levels of these exons. Exons with a low *CV* are good candidates for reference genes, because they exhibit a low variation across all maternal blood samples (Figure 1). For example, those 458 exons (representing 395 genes) detected at a *CV* below the *10^th^ percentile* (*log_10_*(*CV*%) < 1.25) are viable candidates (orange circles in Figure 1; File S1). The range of the *CV* of these 458 exons is 7.75%-17.7%, which are considerably small compared with the *CV* of the least variably detected *GAPDH* exon (27.3%; *log*(*CV*%) = 1.4; Figure 1, black triangle). They were detected at various RNA levels ranging from 1 count to over 16 000 counts. We consider the genes represented by these exons as stably detected in maternal blood.

#### 2.1.3. Pathways and functional annotation terms associated with the genes that are stably detected in maternal blood

To gain insights into the functions of the 359 genes with exons that are stably detected in maternal blood, we statistically tested whether certain pathways or gene ontology (GO) terms [15] that annotate their functions are over-represented using the Protein ANalysis THrough Evolutionary Relationships (PANTHER) classification system [16-18]. Briefly, the percentage of genes associated with each category of pathway or GO terms among the list of stably detected genes was calculated. Then, it was compared with the percentage of genes associated with the corresponding category among the entire list of genes in the reference human genome. The fold of over-representation was reported with a *p*-value after correction for multiple testing by the Bonferroni method. We observed an over-representation of nine cellular component PANTHER GO-Slim terms among our list of stably detected genes, as compared with the reference list of all genes in the human genome (Figure 2A). The 12 over-represented the PANTHER pathways [19] are also shown (Figure 2B). The results of these and six other analyses, including complete GO terms in biological process, molecular function and cellular compartments and the Reactome pathways [20,21] are detailed in File S2.

**Figure 2.**
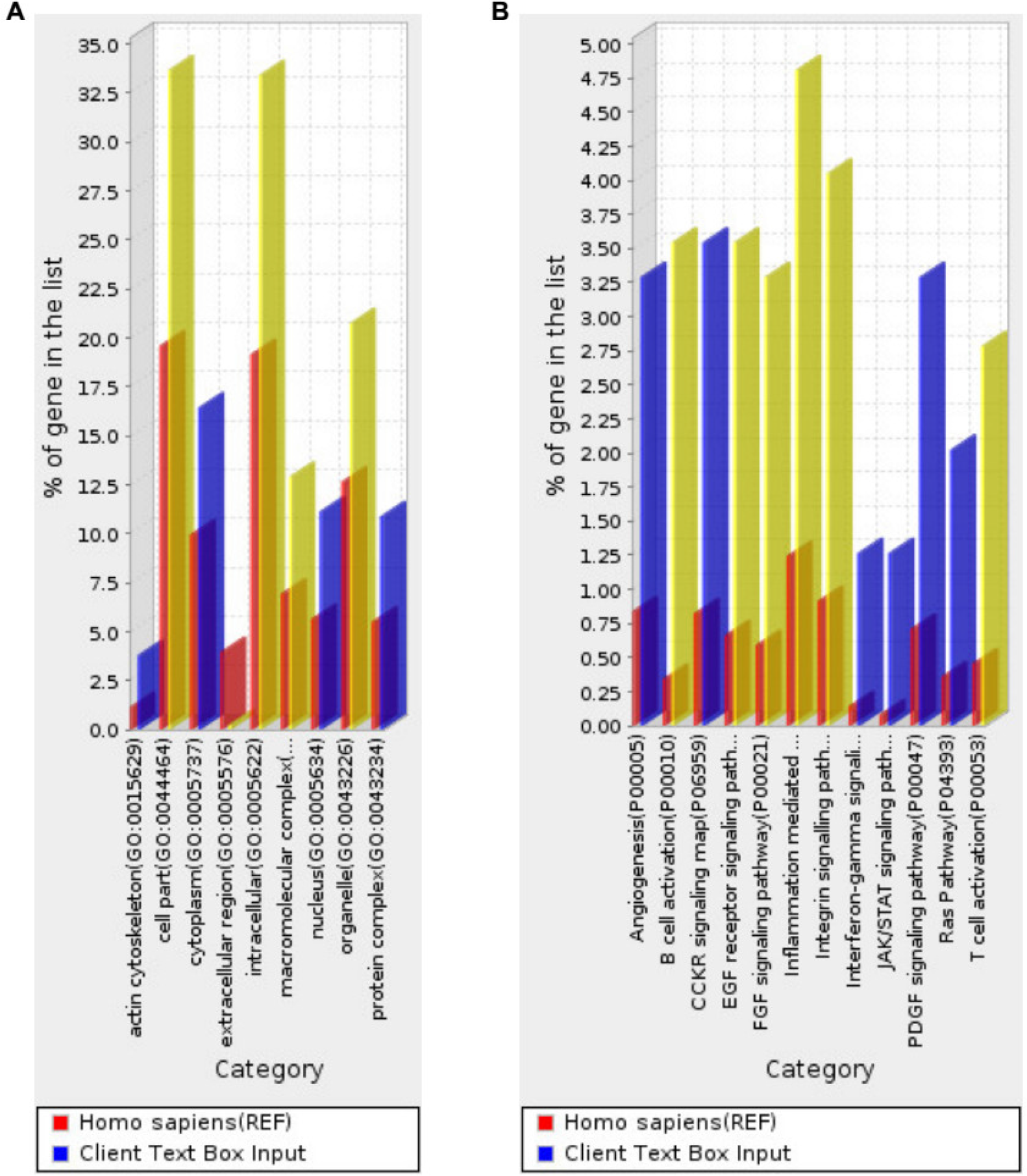
Over-represented annotation terms in the list of 395 genes that are most stably detected in maternal blood. (A) Over-represented PANTHER GO-slim terms in cellular components. (B) Over-represented PANTHER pathways. The list of genes represented by exons detected across all samples at a *CV* in the lowest 10^th^ percentile (orange circles in Figure 1) is subjected to the over-representation test. The percentage of genes in each category of cellular component GO-terms or pathway among this list of most stably detected genes in maternal blood (client input, blue or yellow bars) is shown. It is compared with the percentage of genes in the corresponding category among the entire list of genes in the human reference genome (REF, red bars). *P*-values are adjusted for multiple testing by the Bonferroni method. Blue bars, *p* < 0.05. Yellow bars, *p* < 0.001.

**Figure 3.**
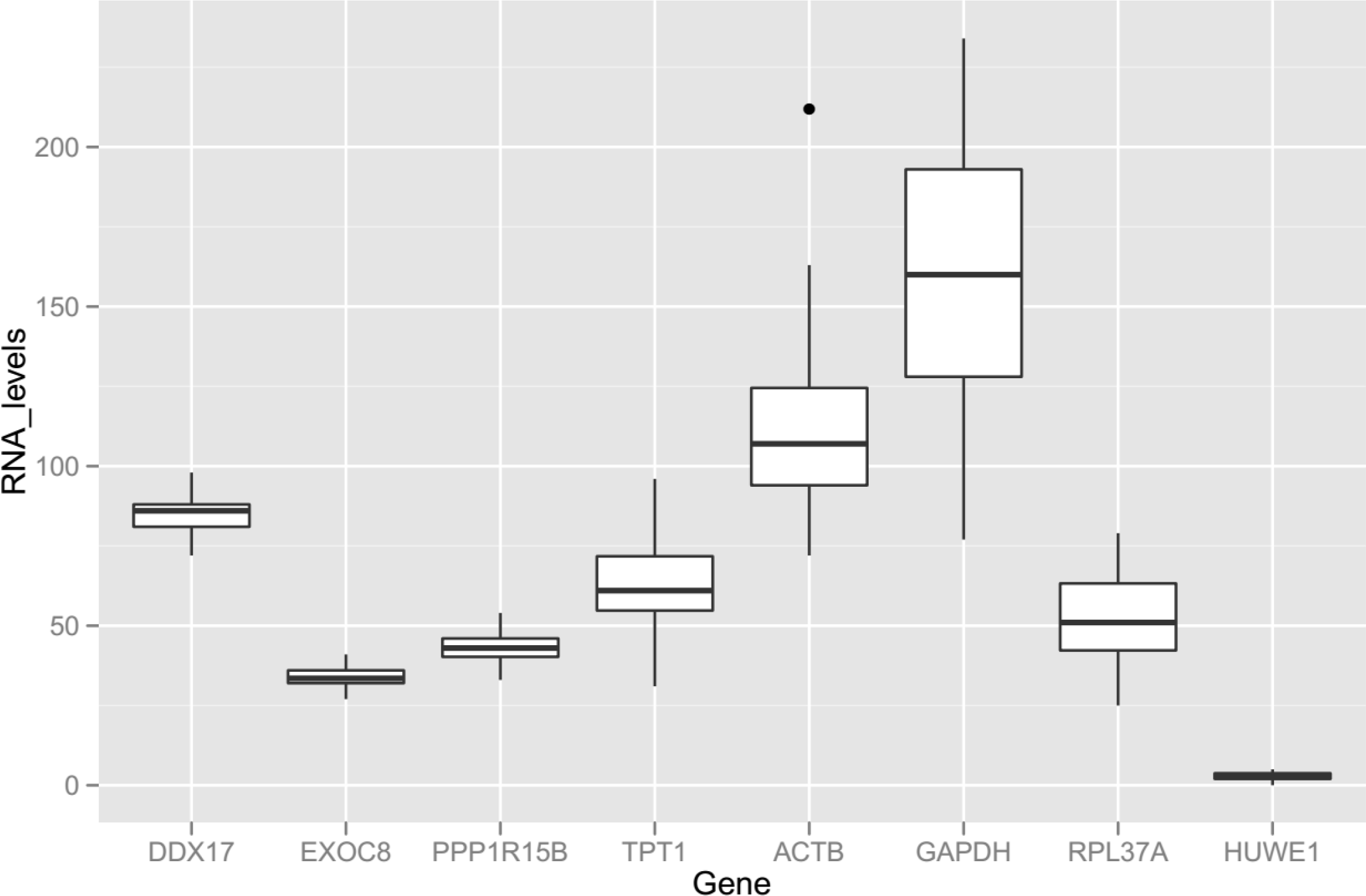
Box plot of RNA levels of eight candidate reference genes in all maternal blood samples subjected to RNA-seq analysis. RNA levels are calculated by normalized read counts, which have taken into account of the varying RNA input, sequencing depths and other technical variations. The bold line inside each box is drawn to the *median*, the bottom and top of each box to the *25^th^* and the *75^th^ percentiles*, the whiskers to the *10^th^* and *90^th^ percentiles*.

Compared with the reference list of all genes in the human genome, the list of 359 genes that are stably detected in maternal blood is over-represented with GO terms in cellular components (Figure 2A; File S2, PANTHER GO-Slim Cellular Component) including macromolecular complex (over-represented by 1.9-fold, adjusted *p*, 9.7×10^-4^), intracellular (1.7-fold; *p*, 9.2×10^-10^), cell part (1.7-fold; *p*, 2.3×10^-9^), organelle (1.6-fold; *p*, 2.7×10^-4^), and actin cytoskeleton (3.3-fold; *p*, 4.1×10^-3^). Overall, this is in line with the major components expected of blood cells after lysis in the RNA extraction step.

Moreover, this list of 359 stably detected genes is over-represented with 12 pathways (Figure 2B; File S2, PANTHER pathways) including B cell activation (10-fold; *p*, 3.0×10^-8^), T cell activation (6.1-fold; *p*, 4.7×10^-4^), inflammation mediated by chemokine and cytokine signaling pathway (3.9-fold; *p*, 1.4×10^-4^), FGF signaling pathway (5.6-fold; *p*, 1.6×10^-4^), EGF receptor signaling pathway (5.4-fold; *p*, 9.7×10^-5^), Integrin signalling pathway (4.4-fold; *p*, 1.8×10^-4^). The former three pathways highlight the important roles played by immune regulation in pregnancy [22], and the latter three mediate growth and proliferation [23-25]. The stable expression or detection of these genes in maternal blood is consistent with pregnancy and a growing fetus.

Furthermore, the list of stably detected genes in blood is over-represented with pathways in other curated database (File S2, Reactome pathways), including regulation of actin dynamics for phagocytic cup formation (6.5-fold; *p*, 2.9×10^-3^), Fc gamma receptor (FCGR) dependent phagocytosis (6.1-fold; *p*, 7.0×10^-4^) and clathrin-mediated endocytosis (4.8-fold; *p*, 1.9×10^-2^). These are consistent with the presence of white blood cells.

#### 2.1.4. Shortlisting of reference genes from our RNA-seq data and the literature

Ideally, the RNA levels of the reference gene and our gene of interest should not differ by more than a few orders of magnitude. We are interested in a panel of 36 preterm birth-associated transcripts that are aberrantly expressed in the preterm placenta and released into maternal plasma [7]. Of these, 12 placental transcripts were observed at a median of 5 counts in our RNA-seq data on maternal whole blood (Table S1). Further, we are interested in preterm birth-associated transcripts that are aberrantly expressed in maternal blood cells of the women presented with preterm labor. The RNA levels of these blood transcripts in whole blood are expected to be higher than those of the placental transcripts. Thus, we shortlisted reference genes that are expressed at RNA levels of about 5 counts to 500 counts and have a low *CV* across maternal blood samples. In our RNA-seq data, the *CV* of RNA levels expressed by the *DDX17* (Ensembl exon ID, ENSE00001942031), *EXOC8* (ENSE00001442235) and *PPP1R15B* (ENSE00001443770) genes are 7.7%, 9.0% and 9.4%, respectively. See Table 1 for the names and symbols of the genes mentioned in this paper. Their *CV* ranked 1^st^, 4^th^ and 6^th^ among the 4 579 robustly detected exons in maternal blood. Their *mean* RNA levels (±*SD*) are 85 counts ± 7 counts, 34 counts ± 3 counts and 27 counts ± 3 counts, respectively.

**Table 1.** Gene symbols and names mentioned in this paper.

| Gene<br>symbol | HGNC<sup>1</sup> approved name | HGNC<br>ID | Location |
| --- | --- | --- | --- |
| *ACTB* | *actin beta* | 132 | 7p22.1 |
| *DDX17* | *DEAD-box helicase 17* | 2740 | 22q13.1 |
| *EXOC8* | *exocyst complex component 8* | 24659 | 1q42.2 |
| *GAPDH* | *glyceraldehyde-3-phosphate dehydrogenase* | 4141 | 12p13.31 |
| *PPP1R15B* | *protein phosphatase 1 regulatory subunit 15B* | 14951 | 1q32.1 |
| *RPL37A* | *ribosomal protein L37a* | 10348 | 2q35 |
| *TPT1* | *tumor protein, translationally-controlled 1* | 12022 | 13q14.13 |
| *HUWE1* | *HECT, UBA and WWE domain containing*<br>*1, E3 ubiquitin protein ligase* | 30892 | Xp11.22 |

<sup>1</sup> HGNC, HUGO Gene Nomenclature Committee. HUGO, Human genome organization.

Additionally, we shortlisted other reference genes for normalizing RT-qPCR data in human blood sample. Previously, other investigators also applied high-throughput data to systematically search for reference genes for normalizing RNA transcripts levels in blood. In a study based on microarray data, among several other genes, *RPL37A* was found to be suitable for normalizing RT-qPCR data in whole blood from tuberculosis patients [5]. *RPL37A* was also robustly detected in our RNA-seq experiment. In a meta-analysis of microarray data on 1 053 blood samples from healthy individuals and patients with cancer and other abnormalities, less than a dozen of genes were found to be expressed at less than a *CV* of 30% [11]. Of these, *ACTB*, *HUWE1* and *TPT1* showed the highest RNA levels. Historically, *GAPDH* was often used as a reference gene, despite the controversy whether its expression is stable enough to serve this purpose [26].

To investigate whether the mentioned genes are suitable for normalizing RT-qPCR data from maternal blood samples, we examined the data distribution of their RNA levels in our RNA-seq data (Figure 2). As expected, the *IQR* of the RNA levels of *DDX17*, *EXOC8*, *PPP1R15B* are small, because they were shortlisted based on their small *CV* in the same dataset. The *IQR* of the RNA levels of *GAPDH* is the largest among the shortlisted genes. The *CV* of the RNA levels for *TPT1* (ENSE00001618156), *ACTB* (ENSE00001902558), *GAPDH* (ENSE00001817977), *RPL37A* (ENSE00001900648) and *HUWE1* (ENSE00000978660) are 21%, 25%, 27%, 29% and 43%, respectively. The highly variable expression levels of the *RPL37A* and the *HUWE1* genes and the low RNA levels expressed by the *HUWE1* gene make them unsuitable to serve as reference genes in maternal blood samples. In fact, of these 6 candidates identified from the literature, the *CV* of 5 genes are larger than the lower quartile of *CV* (24%) as shown by the viable reference gene candidates that may be identified from our RNA-seq dataset (Figure 1, orange circles). Nevertheless, it appears that these candidates from the literature may complement the range of RNA levels not covered by the three candidates shortlisted based on our RNA-seq data. Hence, we embarked on designing RT-qPCR assays to quantify the RNA levels of the *ACTB*, *DDX17*, *EXOC8*, *GAPDH*, *PPP1R15B* and *TPT1* genes in maternal blood samples.

#### 2.1.3. Exon-level RNA-seq data in selected genes

To observe the data distribution of RNA levels of different exons on the same gene, we plotted selected genes with multiple exons detected in our RNA-seq data (Figure 4 and Table 2). We observed much variation in the detection of exons from the same gene in maternal blood, with non-overlapping *IQR* (Figure 4) between them. This observation highlights the advantage of summarizing the RNA-seq data at the exon-level. In designing RT-qPCR assays to quantify the reference gene, it is advisable to target the exon detected at a lower *CV*.

**Figure 4.**
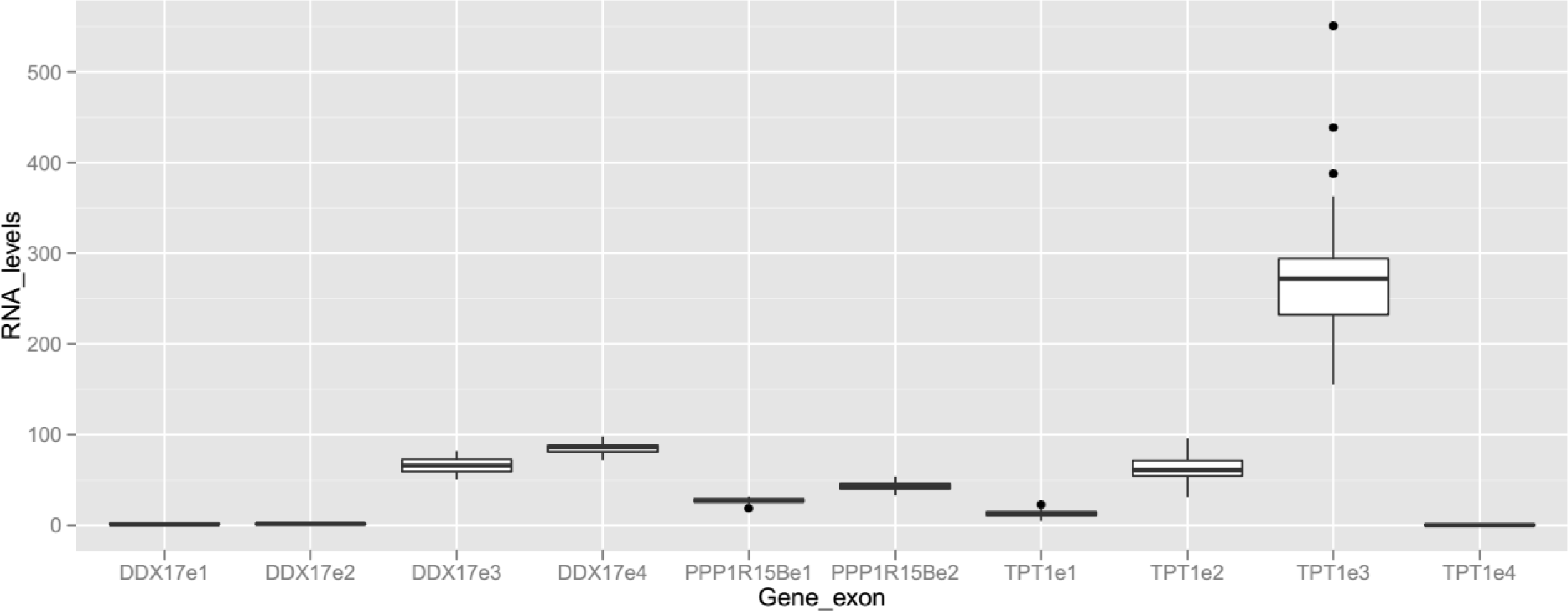
Box plot of RNA levels of multiple exons in selected genes across all blood samples subjected to RNA-seq analysis. Selected exons (e1, e2, etc.) detected in the RNA-seq dataset are shown. See Table 2 for their Ensembl exon ID, *mean, SD* and *CV* of RNA levels. See legend of Figure 3 for the calculation of RNA levels, definition of the box, whiskers and dots.

**Table 2.**
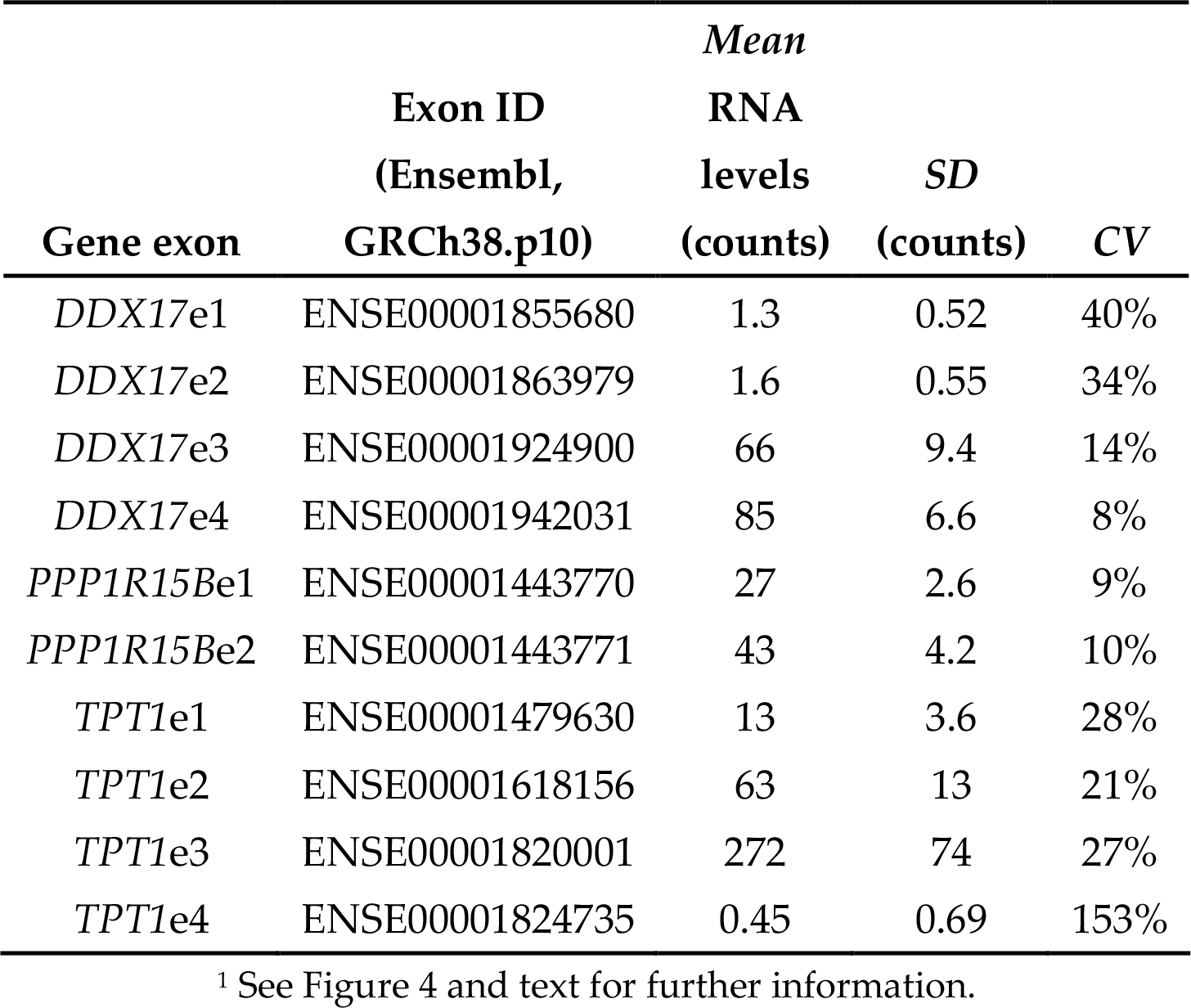
Details of multiple exons in selected^1^ genes detected in the RNA-seq dataset.

The *CV* is large for exons detected at a low mean RNA levels in this RNA experiment (Table 2). For example, *DDX17*e1, *DDX17*e2 and *TPT1*e4. Sequencing deeper than 150 million reads per sample may result in more accurate counting of reads and lower the *CV* of these exons. However, our data suggest that their expression levels are relatively lower than other exons of the same gene. RT-qPCR assays targeting these exons may be less reliable.

### 2.2. Gene expression stability analysis

#### 2.2.1. Comparing complementary algorithms in analyzing gene expression stability

To validate whether the candidate reference genes are stably detected, we assessed gene expression by a different technology platform, namely RT-qPCR, on an independent cohort comprising 32 maternal blood samples which were not used in the RNA-seq experiment. To ensure the best experimental results, the RT-qPCR assays were designed, optimized and performed in compliance with the MIQE guidelines [27]. The experimental details including sequences of the thermal profile, the primer and hydrolysis probe are described in the Methods and File S3.

We used two common softwares, namely geNorm [4] and NormFinder [28], to analyze expression stability of the six selected genes. In the geNorm software, for each candidate reference gene, a stability value (*M*), which is defined as the average pairwise variation of that particular gene compared with all other candidate genes, is calculated. In the NormFinder software, a stability value is also calculated for each gene by a model-based approach, taking into the account of its intra-and inter-group variations. In both softwares, the lowest stability value indicates the most stable gene.

The usual practice is to assess the stability values in a first batch of samples and then presume these values are applicable in other future batches of samples. To test how far this presumption is valid for maternal blood samples collected from women presented with preterm labor, we partitioned the 32 samples into two subsets, each containing approximately equal ratio of sPTB cases to TB controls (Table 3). Then, we calculated the gene stability values by both softwares for the two subsets (Figure 5). The stability values calculated using geNorm in subsets #1 and #2 are correlated (Spearman correlation coefficient, *r* = 0.94, *p* = 0.005), and so are those calculated using NormFinder (Spearman, *r* = 0.89, *p* = 0.02). The 95% confidence interval of the best-fitting line between the two subsets in geNorm appears to be narrower than that in NormFinder (Figure 4, grey zone).

**Table 3.** Samples used for gene expression stability analysis^1^.

|  | Patients<br>ending in<br>sPTB<sup>1</sup> (n) | Patients<br>ending in<br>TB<sup>2</sup> (n) | Total<br>(n) |
| --- | --- | --- | --- |
| Subset #1 | 9 | 8 | 17 |
| Subset #2 | 8 | 7 | 15 |
| Entire set | 17 | 15 | 32 |

**Figure 5.**
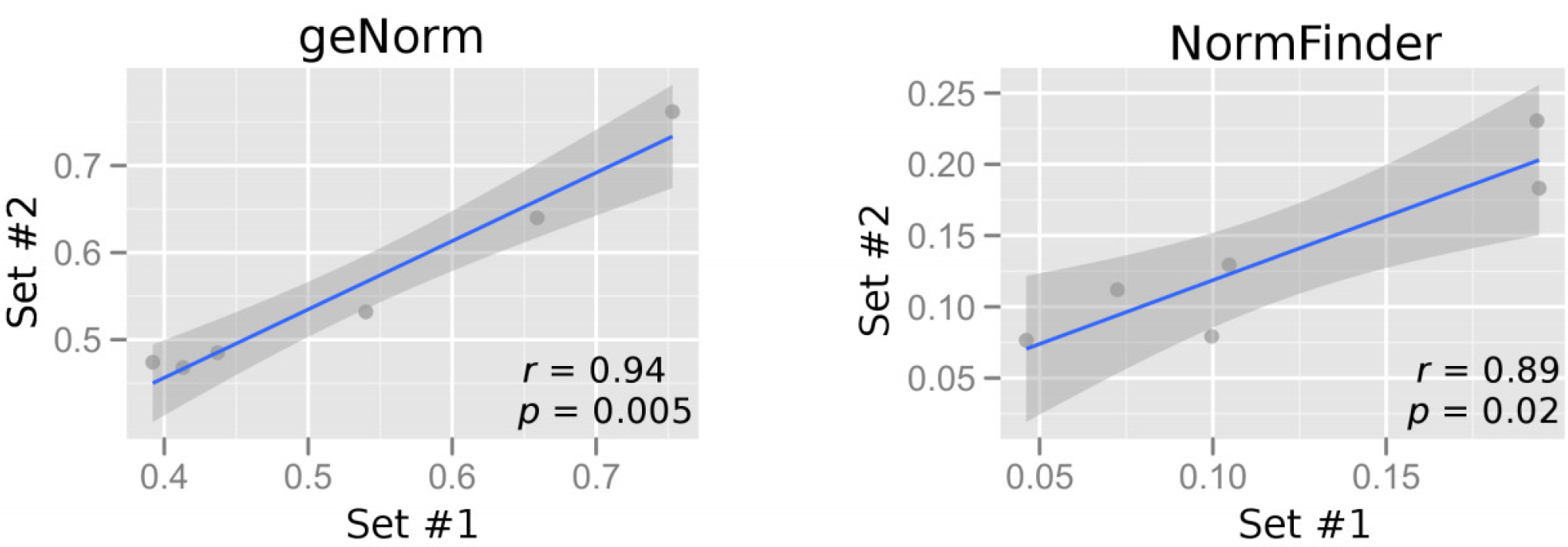
Correlation plots of gene expression stability values from the respective software in sample set #1 *vs* set #2. The geNorm *M* stability value (left) and the NormFinder stability values (right) are plotted. Genes with the smallest stability value have the most stable expression. Each dot represents a gene. The entire set of samples are divided into two subsets, sets #1 and #2, each comprising approximately equal portion of sPTB cases and TB controls. Spearman correlation coefficient *r* (*rho*) and *p* values are shown. Grey zone is the 95% confidence interval of the best-fitting line (blue).

Further, we ranked the candidate reference genes in increasing order of the stability values. In other words, the most stable gene ranks first. The ranks for each gene in the two subsets and the absolute rank difference between them were tabulated (Table 4). The sum of the rank differences for all six tested gene is smaller for the stability values calculated by the geNorm software than those by the NormFinder software. Taken altogether, for the maternal blood samples and the candidate reference genes that we tested, geNorm gave slightly more reproducible results between the two subsets, compared with NormFinder.

**Table 4.**
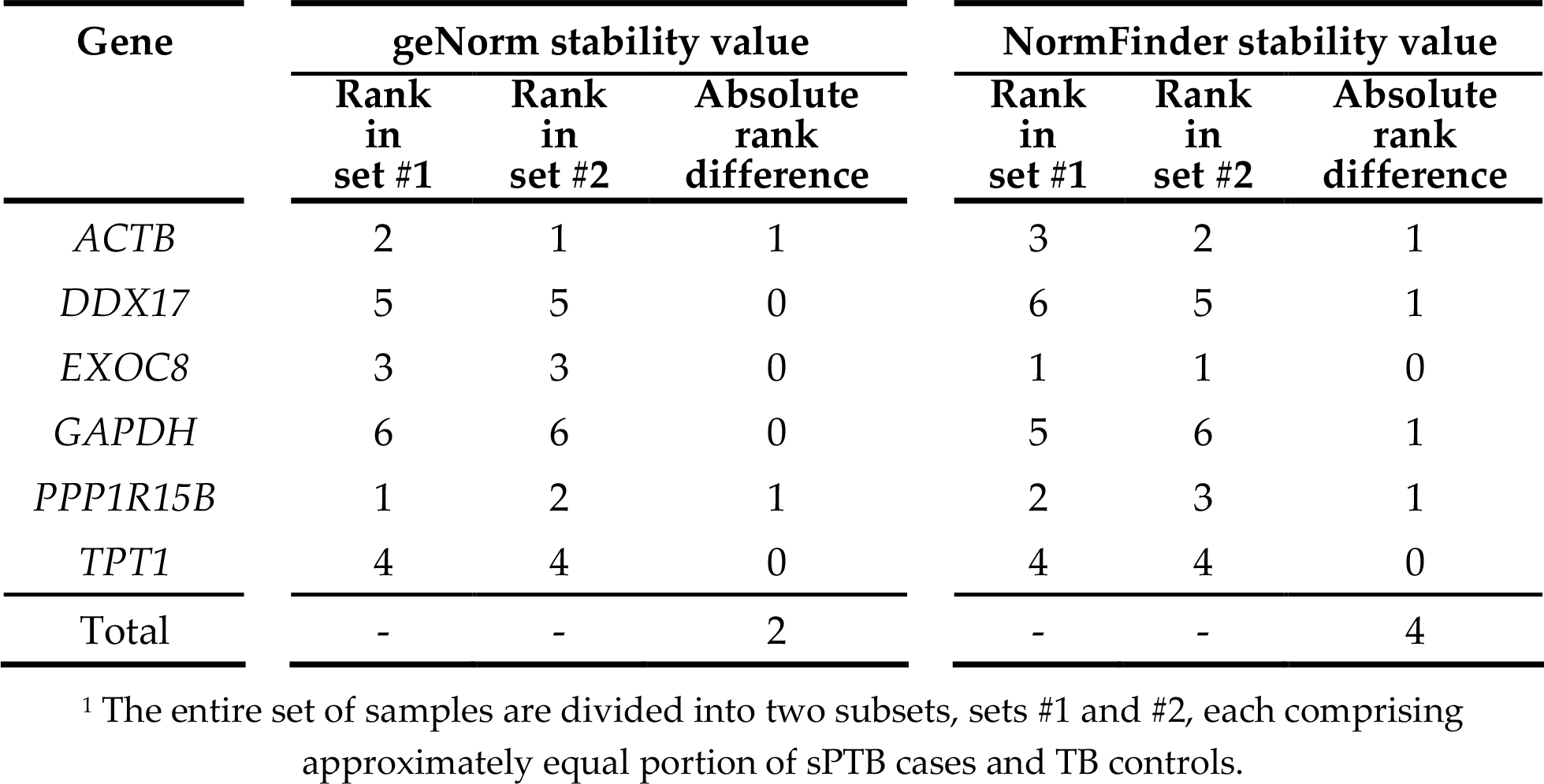
Ranking of stability values calculated by the respective software in two sets of samples^1^.

| Gene | geNorm stability value |  |  | NormFinder stability value |  |  |
| --- | --- | --- | --- | --- | --- | --- |
|  | Rank in set #1 | Rank in set #2 | Absolute rank difference | Rank in set #1 | Rank in set #2 | Absolute rank difference |
| *ACTB* | 2 | 1 | 1 | 3 | 2 | 1 |
| *DDX17* | 5 | 5 | 0 | 6 | 5 | 1 |
| *EXOC8* | 3 | 3 | 0 | 1 | 1 | 0 |
| *GAPDH* | 6 | 6 | 0 | 5 | 6 | 1 |
| *PPP1R15B* | 1 | 2 | 1 | 2 | 3 | 1 |
| *TPT1* | 4 | 4 | 0 | 4 | 4 | 0 |
| Total | - | - | 2 | - | - | 4 |

<sup>1</sup> The entire set of samples are divided into two subsets, sets #1 and #2, each comprising approximately equal portion of sPTB cases and TB controls.

#### 2.2.2. Assessing gene expression stability in maternal blood samples

Because the geNorm software gave the slightly more reproducible results, we proceeded to use it to analyze the entire set of 32 maternal blood samples. Based on stepwise exclusion of the least stable reference gene using the geNorm software, the stability values of the remaining control genes are plotted and the genes are ranked (Figure 5). The candidates in descending order of gene expression stability are *PPP1R15B*, *ACTB*, *EXOC8*, *TPT1*, *DDX17* and *GAPDH*.

Normalization factor (*NF_n_*) of a certain number (*n*) of genes was calculated, starting with the most stable genes. To determine the possible need or utility of including a certain number of genes for normalization, the pairwise variation *V*_*n*/*n*+1_ was calculate between *NF _n_* and *NF* _*n*+1_ (Figure 6). Based on the data in the geNorm paper, if *V*_*n*/*n*+1_ > 0.15, the added gene has a significant effect and should preferably be included for calculation of a reliable *NF* [4]. In our gene stability analysis, the *V*_2/3_ was 0.15. Thus, adding the third most stable reference gene has little effect on the *NF*_3_, compared with *NF*_2_. Hence, the optimal number of reference genes is 2. As such, the optimal *NF* can be calculated as the geometric mean of *PPP1R15B* and *ACTB*.

**Figure 6.**
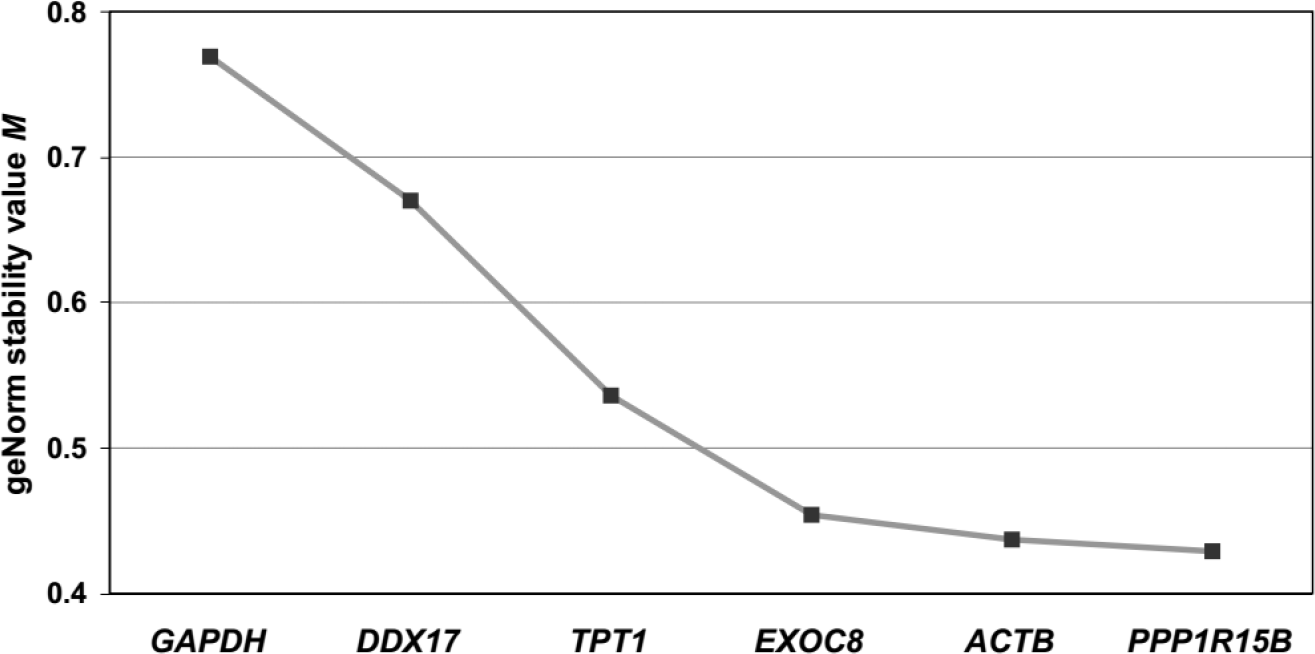
Average expression stability values (*M*) in the entire set of maternal blood samples. *M* is calculated using geNorm of the remaining control genes during stepwise exclusion of the least stable control gene. Gene with the lowest *M* has the most stable expression.

**Figure 7.**
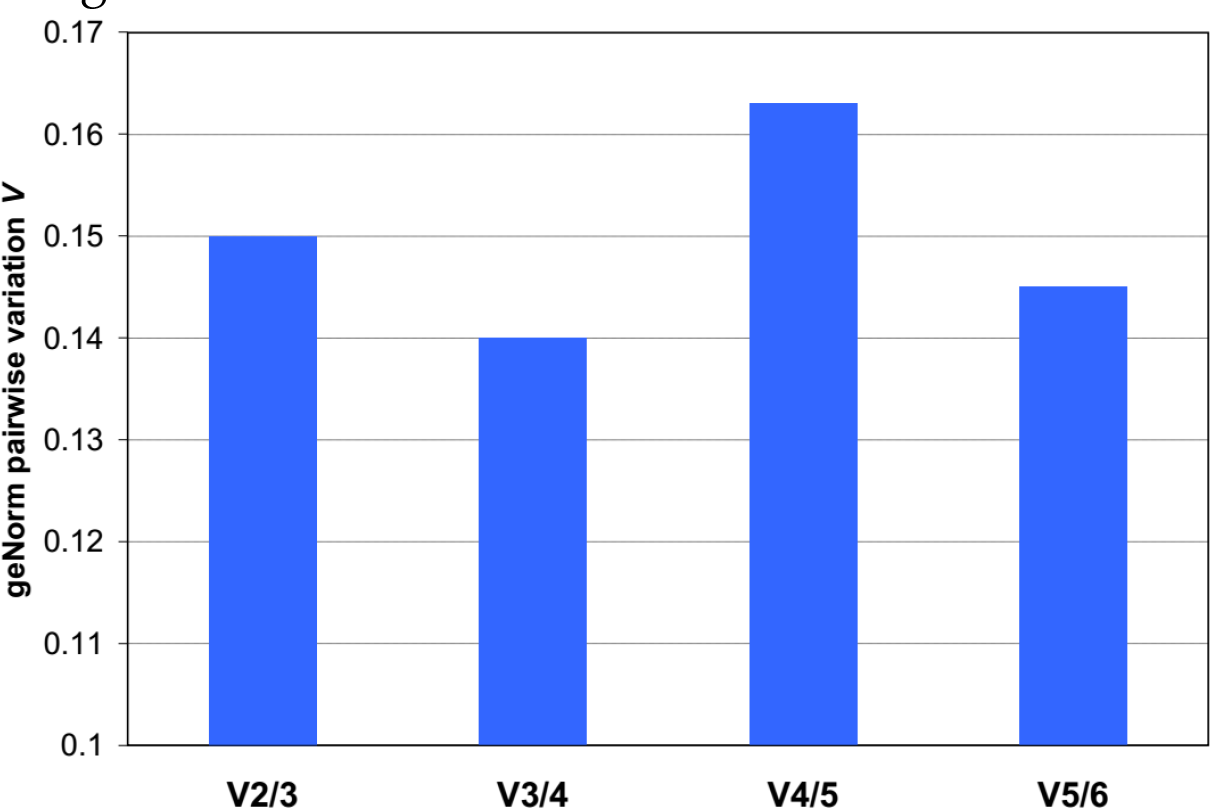
Pairwise variation analysis between normalization factors in the entire set of maternal blood samples. To determine the optimal number of reference genes required for accurate normalization, we perform the pairwise variation (*Vn/n+1*) analysis between the normalization factors *NFn* and *NFn+1*, where *n* is the number of reference genes.

## 3. Discussion

RNA-seq has facilitated us to search for candidate reference genes from the whole-transcriptome. Compared with gene expression microarray, RNA-seq is not limited by a fixed number of probes and known gene sequences. Consequently, considerably more reference gene candidates expressed were identified in this RNA-seq study (Figure 1) than a similar microarray study [11]. However, because the amount of RNA that could be extracted from this preciously collected cohort is limited, it is impractical to systematically validate all candidates using RT-qPCR. Thus, we have validated candidates which are expressed at similar levels as our genes of interest.

RNA-seq, which was performed at a reasonable sequencing depth, has enabled us to profile the transcriptome at higher resolution of the exon level. As illustrated by the three selected genes in Figure 4, the exons of the same gene are detected at considerably different levels. This implies that exon-level data may guide the design of RT-qPCR assays to more specifically target a sub-region that is advantageous for the quantification of a gene transcript. In the case of designing RT-qPCR assays for reference genes, the exons with the lowest *CV* are the preferred targets.

We have also been able to assess the variations of candidate reference genes in terms of *CV* and *IQR* of the normalized read counts on the RNA-seq data. Still, there are limitations of such assessment. The total RNA used for constructing the sequencing library contains a considerable portion of ribosomal RNA (rRNA). Its presence may affect the accuracy of quantification of the amount of RNA input for library construction. Moreover, since the predominance of rRNA often masks the signals from other more informative mRNA in RNA-seq experiments, a common practice is to reduce rRNA before library construction. However, such practice contributes to a source of technical variations. Combining with the quality-filtering, mapping and normalization steps in data analysis, each RNA-seq dataset contains a considerable amount of noise, which should be vigorously controlled for reliable gene expression profiling. Therefore, it is important to validate the findings from RNA-seq by independent technology platform, such as RT-qPCR, and cohort. This is especially true for identification of reference genes, because the impact of the technical variation in RNA-seq on gene expression stability analysis has not been extensively studied. Further, we have supplemented our RNA-seq based selection of candidates with relevant studies which employed other platform for profiling transcriptome.

We have used two complementary approaches for validating the expression stability of the selected reference genes. Both geNorm and NormFinder gave highly correlated (Spearman, *r* = 0.94 and 0.89, respectively) stability values between two different subsets of samples. Thus, the data suggest that both softwares generate similar stability values across different batches of samples. Although geNorm has generated slightly more correlated stability values across the two subsets, it is noted that both geNorm and NormFinder have consistently ranked *PPP1R15B, EXOC8* and *ACTB* as the most stably detected genes (Table 4). Similarly, in the analysis of the entire set of 32 samples, geNorm has also ranked the same three genes as the most stably detected genes (Figure 6). Apparently, for whole blood samples collected from women during the presentation of preterm labor, *PPP1R15B, EXOC8* and *ACTB* are suitable to serve as reference gene for normalizing the RNA levels of other circulating RNA transcripts. On a related note, among the genes selected for expression stability analysis, *GAPDH* was essentially consistently ranked as the least stable by both softwares in both subsets (Table 4) and the entire set (Figure 6). Our data suggest that reference genes with lower variation than *GAPDH* do exist in maternal blood.

We observed that the RNA-seq data is not always predictive of the gene expression stability analysis data based on the RT-qPCR. For instance, in the RNA-seq data, the RNA levels of *ACTB* were detected at a *CV* of 25%, which is more variable than those of *DDX17* (7.7%), *EXOC8* (9.0%) and *PPP1R15B* (9.4%). On the other hand, in the geNorm analysis, *ACTB* turned out to be one of the most stably detected genes. A closer examination revealed that the *ACTB* exons targeted by the commercially available RT-qPCR assay were not detected in our RNA-seq dataset. The RT-qPCR assay pre-designed by that company targeted an amplicon that is mapped to more than one genomic location. This type of exonic sequencing reads been filtered out after mapping, because they may interfere with the counting of transcripts in the RNA-seq experiment. This illustrates how a search for reference genes based on RNA-seq may benefit from the relevant studies using an alternative technology.

The major advantage of our RNA-seq approach is that reference gene candidates could be systematically identified from the whole-transcriptome, which is more comprehensive. Moreover, the exon-level data allow a more specific design of RT-qPCR assays. The downside is that certain short sequencing reads that are mapped to more than one unique location in the genome are missed. For instance, the *ACTB* exon targeted by the commercial RT-qPCR assay was not detected in our RNA-seq data. It has been estimated that the RNA levels of hundreds of genes in the human genome could not be quantified accurately by RNA-Seq [29]. These genes are enriched for gene families, and many of them have been implicated in human disease. Usually, these ambiguously-mapped reads are discarded to improve the accuracy of quantification the uniquely-mapped reads in RNA-seq analysis. Recently, methods for including these non-unique reads in RNA-seq analysis have emerged. For instance, it is now possible to quantify the RNA levels of a family of similar sequences as a group [29]. Combined with the increasingly longer read length facilitated by newer sequencing platforms and reagents, we expect this negative impact of ambiguous reads on RNA-seq analysis will be minimized.

In this study, we provided a list of 458 exons (395 genes) that are most stably detected in maternal blood. Unlike many other tissues, peripheral blood is readily and non-invasively obtainable from a patient. Many RNA transcripts expressed in other tissues are often released into the blood circulation, including preterm birth-associated placental RNA and pregnancy-associated microRNA [7, 30]. Nevertheless, the potential use of these circulating RNA as biomarkers is hindered by the paucity of study on reference genes in peripheral blood. Shortlisting from the transcriptome data and validating them by RT-qPCR are two essential but resource-demanding steps in finding suitable reference genes. Although the reference genes suitable for normalizing blood expression data from patients presented with preterm labor may not be suitable for data from patients suffering from other conditions, our list of stably detected exons (Figure 1) could still be useful for normalization of other experimental systems.

To find reference genes in blood samples from patients with other diseases, one may first shortlist about 10 stably detected exons from our RNA-seq data (File S1) with similar RNA levels to the gene of interest. Then, RT-qPCR assays targeting the specific exons could be designed for gene expression stability analysis by geNorm, NormFinder or the like. Thus, the time and resources to perform the whole-transcriptome experiment and bioinformatics analysis can be saved. Since blood cells are the predominant source of nucleic acids in cell-free plasma [31], we reason that our RNA-seq data on whole blood will also serve as a starting point for finding reference genes in plasma.

In addition, we noted that the stably detected genes in maternal blood are over-represented with annotation terms in macromolecular complex and actin cytoskeleton (Figure 2A). This is not surprising, because they are the major structural components of any cells. Interestingly, other terms are associated with phagocytosis (File S2, Reactome pathways), B cell activation, T cells activation and inflammation (Figure 2B). This is consistent with the notions that pregnancy and preterm birth are associated with immune regulation and infection, respectively.

Intriguingly, terms in FGF, EGFR signaling pathways and Integrin pathways are also over-represented in the stably detected genes. Members of the FGF family function in the earliest stages of embryonic development and during organogenesis to maintain progenitor cells and mediate their growth, differentiation, survival and patterning [23]. The EGFR signaling pathway is one of the most important pathways that regulate growth, survival, proliferation, and differentiation in mammalian cells [24]. Integrins contribute to cell growth by providing a physical linkage between cytoskeletal structures and the extracellular matrix, and also by participating in various signal transduction processes [25]. The stable detection of genes in these pathways and fetal growth warrants further investigation. Will these genes remain stably detected in maternal blood of pregnancy complicated by intrauterine growth restriction, macrosomia or gestational diabetes?

Taken altogether, the maternal circulation harbors RNA transcripts that are stably detected. We identified a list of 395 genes that were detected at a low *CV* in maternal whole blood samples collected from women presented with preterm labor. They are good reference gene candidates for normalizing expression data from these patients. Our list may also be useful as a starting point for finding reference genes for normalizing data from patients with other disease or from samples of different blood compartments. With reference genes that are more stable, it is hoped that the hurdles of normalization could be overcome and that more differentially expressed transcripts could be identified in the circulatory system.

## 4. Materials and Methods

### 4.1. Recruitment of study subjects

This study was approved by the institutional review boards (CRE 2012.032 and 2013.10-85). Women presented with preterm labor and fulfilled the inclusion and exclusion criteria were invited to participate in this study. The inclusion criteria are women with (i) uterine contractions at least once every 10 minutes < 34 weeks, (ii) intact membrane, (iii) singleton pregnancies, and (iv) a Chinese or Korean ethnicity. The exclusion criteria are women with pregnancies complicated with (i) preterm prelabor rupture of membrane, (ii) multiple gestation, (iii) preeclampsia, (iv) fetal growth restriction, (v) macrosomia, (vi) fetal distress, (vii) antepartum hemorrhage, (viii) fetal chromosomal or structural abnormalities, (ix) history of uterine abnormality or cervical surgery, and (x) indicated preterm births before 37 weeks (induction of labor, elective or emergency term cesarean deliveries), where deliveries are iatrogenic. Gestational ages were established based on menstrual date confirmed by sonographic examination < 20 weeks. We followed up their delivery outcome and categorized them accordingly into the spontaneous preterm birth (sPTB, before 37 weeks of gestation) and the term birth (TB, on or after 37 weeks) groups.

### 4.2. Blood collection and RNA extraction

Blood sample (9 mL) was collected from the antecubital fossa in EDTA-containing tubes (Greiner Bio-One). To minimize RNA degradation, the blood sample was mixed with RNAlater (Thermo Fisher Scientific) shortly after venesection and stored at -80°C until extraction. RNA was extracted using the RiboPure RNA Purification Kit (blood) (Thermo Fisher Scientific) and treated with *DNase* I (Thermo Fisher Scientific) to minimize contaminating genomic DNA. Quality of the RNA preparation was assessed by spectrophotometry and RNA Pico chip on the Bioanalyzer (Agilent).

### 4.3. RNA-seq

Forty libraries (2 technical replicates per blood sample) were constructed for strand-specific pair-end cDNA sequencing (TruSeq 3000 4000 SBS Kit v3) according to the TruSeq Stranded Total RNA Sample Prep Guide (Part # 15031048 Rev. E). To minimize the highly abundant but uninformative ribosomal RNA and globin mRNA transcripts from masking the signals from the more informative transcripts, we subjected the RNA samples to pre-treatment by Ribo-Zero Globin (Illumina). RNA-seq was performed on the HiSeq 4000 sequencer (HiSeq 3000 4000 System User Guide Part #15066496 Rev. A HCS 3.3.30) using control software HCS v3.3.20. We filtered out low-quality sequences, trimmed away adapter sequences (Trimmomatic, v.0.33) [12]. To remove technical variations, we performed normalization using the R-package RUVSeq (ver. 1.6.0) according to instructions in the manual compiled May 3, 2016 [14]. The RUVs method was used to estimate the unwanted variation using replicate samples. We calculated the mean and *CV* of RNA levels at the exon level. When multiple exons were robustly detected in the RNA-seq dataset, the data from the exon with the lowest *CV* were reported unless otherwise stated.

### 4.3. RT-qPCR

To leverage on the exon-level transcriptome data, we designed RT-qPCR assays to target the exon with the lowest *CV* of RNA levels for all except two genes in this study using the PrimerQuest software (Integrated DNA Technologies, IDT). The assays for *ACTB* and *GAPDH* genes were predesigned by and purchased from Integrated DNA Technologies, Inc. To improve the accuracy of quantification, hydrolysis probes were used. Primers and probes were validated *in silco* for possible secondary structures and non-specific binding by Primer-BLAST [32]. The full probe and primer sequences, reaction conditions and PCR efficiencies are shown in File S3. To avoid contaminating genomic DNA, we pre-treated the extracted RNA samples with *DNase* I before RT-qPCR. To monitor for environmental contamination, no template controls were run in parallel.

A RT-qPCR primed by random primers is linear over a narrower range than a similar reaction primed by target-specific primers [33]. Further, it was shown that use of random hexameric primer sequences for reverse transcription may overstate the actual amount of mRNA up to 19 times [34]. Therefore, for more accurate results, we used gene-specific primers for the reverse transcription step in RT-qPCR. To minimize technical variation and the risk of contamination, we performed one-step RT-qPCR using the RNA Master Hydrolysis Probe kit (Roche), which involves no opening of the reaction vessel after addition of the RNA template. For the gene expression stability analysis, 10 ng of *DNase* I-treated RNA was added to each reaction. The reactions were performed on the LC480 platform (Roche). We performed the geNorm analysis in qBase PLUS, which is MIQE-compliant [35]. PCR efficiency of each assay was determined by a calibration curve constructed from standards of known concentrations.

### 4.3. Other data analyses

Unless mentioned above, data analysis including correlation tests, descriptive statistics and graph plotting were performed using DeduceR [36] which is a graphical user interface for R and Microsoft Excel.

## Supplementary Materials

File S1. Details of exons detected with a *CV* within the *10^th^ percentile* among 4 579 exons robustly detected in maternal blood.

File S2. Pathways and Gene Ontology terms associated with 359 stably detected exons in maternal blood.

File S3. Technical details of RT-qPCR assays used in this study.

Table S1. RNA levels of transcripts that are aberrantly expressed in the preterm placenta and released into maternal circulation.

## Acknowledgments

This work was supported by the Health and Medical Research Fund of the Food and Health Bureau of the Hong Kong SAR Government (Project Ref No.: 01120066).

## Author Contributions

S.S.C.C., T.-F.C. and T.-Y.L. conceived and designed the experiments; K.K.W.W. and S.K.W.L performed the experiments; C.Y.L.C., J.S.L.K., S.S.C.C., T.-F.C. and S.K.W.T analyzed the RNA-seq data. K.K.W.W., S.K.W.L. and S.S.C.C. analyzed the RT-qPCR data; T.-Y.L., K.-Y.L., C.-Y.L., Y.K.Y.C., A.S.Y.H., M.M., O.-K.C. contributed research samples and clinical data; S.K.W.T., T.-F.C. contributed analysis tools; S.S.C.C. wrote the paper.

## Abbreviations

*ssRNA-seq*: Strand-specific RNA sequencing
*RT*: Reverse transcription
*qPCR*: Quantitative polymerse chain reaction
*sPTB*: Spontaneous preterm birth
*TB*: Term birth
*CV*: Coefficient of variation

